# Acoustic cues and season affect mobbing responses in a bird community

**DOI:** 10.1101/2022.05.05.490715

**Authors:** Ambre Salis, Jean-Paul Léna, Thierry Lengagne

## Abstract

Heterospecific communication is common for birds when mobbing a predator. However, joining the mob should depend on the number of callers already enrolled, as larger mobs imply lower individual risks for the newcomer. In addition, some ‘community informant’ species seem more reliable regarding the information transferred in mobbing calls. Birds should therefore rely on both the number of callers and the species identity of the caller(s) when mobbing. In the present study, we tested the potential interaction between two acoustic cues. In a playback experiment, we modified the number of callers (through an increased number of calling individuals correlated to an increased duty cycle) and the emitter species (crested tits versus coal tits). Overall, we found that soundtracks with three callers triggered more mobbing than soundtracks with one caller and that soundtracks with coal tits’ calls triggered more mobbing than soundtracks with crested tits’ calls. Our results therefore support the hypothesis that birds consider both the species and the number of callers when joining a mobbing chorus in winter. Finally, we replicated the experiment in spring and did not record the same responses from the bird community. Indeed, only soundtracks with three coal tits triggered a mobbing response, suggesting therefore that the seasonal context can affect the results of studies on heterospecific communication. The potential mechanisms implicated in the varying responses to different acoustic cues and different seasons are discussed and should deserve further investigations.

## INTRODUCTION

Clustering around a predator and actively harassing it instead of fleeing is a widespread phenomenon termed ‘mobbing’. Particularly common in birds (Carlson et al. 2018), mobbing encourages the predator to give up hunting and move to another location in both the short and long term (the Move-On Hypothesis, Curio 1978, Flasskamp 1994). Other benefits, such as monitoring the predators and enhancing learning opportunities for offspring, have been proposed (Curio 1978). Costs associated with such behavior are however non-negligible: in addition to the loss of time and energy when responding to an individual calling, the direct confrontation with a predator could result in direct aggression from the predator (Curio and Regelmann 1986, Poian and Yorke 1989, Sordahl 1990). Mobbing efficiency (i.e., the ratio of costs / benefits) can be improved by increasing the number of mobbing individuals (Krams et al. 2010, Wheatcroft and Price 2018). Indeed, larger groups decrease both the individual risk of being targeted by the predator (Hamilton selfish herd or dilution effect, Foster and Treherne 1981), and the overall success of the predator through confusion effect (Carlson et al. 2018). Larger groups also increase the chances of repelling the predator (Hendrichsen et al. 2006). Such an increase of participants can be achieved both with conspecific and heterospecific individuals, and heterospecific mobs are indeed well documented (e.g., Dutour et al. 2017a, Goodale and Kotagama 2005, Hua et al. 2016). Although heterospecific mobbing responses probably emerged as simple by-product mutualism (Kostan 2002), the relationships between species can be complex. Indeed, participation in such mobs is often unequal (Dutour et al. 2017b), with some species risking less by following the group at a distance (Magrath et al. 2015). In opposition, other species seem particularly active and trustworthy regarding the information conveyed in the calls (Farine et al. 2015). For such species that are active, reliable, and highly responded to, the term ‘community informant’ has been proposed (Carlson et al. 2020).

The rationale to join mobbing birds should therefore depend on two main acoustic cues: (i) the number of birds already mobbing, as a greater number of birds indicates a lower risk for new participants, and (ii) the species identity of the caller(s), since some species convey more reliable and relevant information than others. To test these hypotheses, we built a set of playback experiments using a factorial design. We broadcast soundtracks of either one or three coal tits (*Periparus ater*) and one or three crested tits (*Lophophanes cristatus*) to free-ranging birds of both species, and recorded their behavioral response (calling and approaching, the most conspicuous signs of mobbing in birds). Following a recent study (Carlson et al. 2020), coal tits and crested tits contrast in their call reliability (i.e., coal tits vary their calls when facing different threats) and heterospecific attraction when mobbing a predator.

Heterospecific communication related to mobbing is prevalent in winter in passerines communities (Dutour et al. 2019), notably because of an increased tendency to flock with heterospecifics to increase predator defense and foraging efficiency (Goodale et al. 2015). We therefore chose to test first and foremost birds during winter. Yet, we also replicated the same experiment in spring to test whether seasonal context could influence experiments about heterospecific communication. Indeed, throughout the year, the physical and social environment of birds varies greatly, possibly impacting their communication (e.g., Clucas et al. 2004, Jiang et al. 2020). In spring, the increased aggressiveness due to territoriality and nest defense could affect results on mobbing behavior (Betts et al. 2005, Jiang et al. 2020). By replicating this experiment in a different season, we test how environmental parameters such as season can affect our biological conclusions.

Our experiment therefore aims at determining the relative flexibility of heterospecific relationships and stability of response to acoustic cues throughout birds’ seasonal activity. By looking at the mobbing response of both coal and crested tits to each other’s calls, as well as the mobbing response of the overall community, we aim at determining how context affects the acoustic cues used by birds when investing in mobbing.

## MATERIAL AND METHODS

### Study site & Species

The playback experiments described below were all done in the Haut-Bugey region, France. This region is a small mountain environment (altitude: ~ 800 m), with mixed deciduous-coniferous forests. Densities of coal and crested tits are high in this area, as shown by the long-term ornithological census in the region: both species were detected in 94% of points, spaced at 150 m from each other (participative database Faune-ain.org administered by the LPO AuRA DT Ain). In this region, small birds are often predated by several predator including the Eurasian pygmy owl *Glaucidium passerinum*. Previous experiments in the region have shown a mobbing response from a large number of species, including the coal and the crested tits (Dutour et al. 2016, Dutour et al. 2017b). When mobbing occurs, birds approach the predator cue and produce calls often with specific aggressive postures (e.g., wing flicks and frequent hops), but direct attack is rare (Carlson & Griesser, 2022).

### General organization

We aimed at testing the mobbing response of free-ranging birds to different soundtracks. To this aim, we established 100 spots for the playback tests in a 10 km^2^ area of coniferous forest in the East of France (46°13’05.0”N 5°41’50.8”E). Each spot was selected along an existing trail but close to a tree allowing birds’ approach and concealment of experimenters. All spots were separated by ~ 100 m (mean and standard deviation: 110.9 ± 27.2 m) since this distance is sufficient to degrade bird sounds (Morton 1975). In addition, we performed a complementary subset of experiments (n = 22 birds tested, 9 crested tits and 13 coal tits) to verify that birds do not follow the observer between successive spots. For this purpose, we followed the same methodology than the one used by Salis et al. on great tits (2022). More specifically, both observers were equipped with the acoustic material and binoculars, and after each test, while one observer was launching the playback experiments on a subsequent location, the other was following the birds from the previous location. We found that from one test to the next one, no bird followed us, and no bird moved farther than 50 m from their original position (see details in Supplementary File 1). While birds can travel large distances in a short period, it is unlikely that we tested the same birds in consecutive tests in the present experiment given the absence of human following and the absence of attraction from the subsequent playbacks.

We created a factorial design in which our four different treatments (different emitter species and number of callers, see paragraph Playbacks for details) were broadcast on each spot. These experiments were first carried out in winter, and then replicated in spring. Each spot consequently received eight playback tests. We avoided spatial and temporal autocorrelation by (i) alternating the four treatments at consecutive spots, and (ii) doing the same number of tests of each treatment, each day. The 400 tests in each season were done in a short period (two weeks) to avoid a potential intra-seasonal effect, and each consecutive test spaced by at least five minutes (each consecutive test was at a different spot, so that each spot was tested only once per day). We changed the order in which the spots were tested each day (different beginning point each day and different directions in the trails). Post hoc analyses (Sup. File 2) show no effect of order of playback treatment nor of the repeated presentation of playbacks on our results.

### Playbacks

We created four treatments: soundtrack with only one calling coal tit (1CO), three coal tits calling simultaneously (3CO), only one calling crested tit (1CR), and lastly, three crested tits (3CR). We did not use a negative control (e.g., heterospecific song or background noise) since we were interested in the difference between our treatments. Moreover, background noise has been used in several studies (Dutour et al. 2019, Salis et al. 2022, Suzuki et al. 2016) and never triggered a response from Parids. To prepare our soundtracks, we elicited mobbing calls from wild crested tit and coal tit by broadcasting a mobbing chorus of various birds (including coal and crested tits, Dutour et al. 2016). Once birds arrived to mob, they were recorded with a ME-67 Sennheiser microphone connected to a K6 basis and a Fostex FR2LE recorder (recording distance of 5 m to 15 m). At last, the recordings were then cleared of any other bird call, their amplitude homogenized at 50% on the entire file with AvisoftSasLab (Avisoft Bioacoustics, Glienicke, Germany), and saved as WAV files. We selected recordings with a number of calls around the mean (± 1 SD) of previous recordings obtained by our team (coal tit: 82 ± 26 notes per min, *N* = 30, crested tit: 134 ± 44 notes per min, *N* = 10). For the treatments with three birds (trio treatments), we superimposed recordings of three different birds calling to simulate a chorus. As a result, the final duty cycle (i.e., the amount of signal present in the playbacks) was higher for the three-birds treatments (~ 9 sec) than for the one-bird treatments (~ 6.5 sec, details in Sup. File. 3). Nevertheless, the calls substantially overlapped, reducing the risk for the focal birds to consider the three-birds treatments as only one bird calling intensely. For each treatment, we built five different soundtracks to circumvent the idiosyncrasy of recorded subjects (Kroodsma 1989).

### Test procedure

One test consisted in playing 30 sec of a mobbing call sequence at each spot with a Bose Soundlink Revolve loudspeaker perched on a tripod (H: 1 m), put near a tree and at an amplitude of 84.01 ± 2.70 dB (calculated at 1 m with Lutron SL-4001, C weighting, slow settings, re. 20 μPa, Templeton et al. 2016). 30 sec is enough to trigger a mobbing response from nearby birds (previous recordings were obtained with such a stimulation), who can approach and call as a response, sometimes with additional aggressive behavior (e.g., wing flicking, Salis et al. 2021). A stimulation of only 30 sec also limited the influence of the first birds to call on the following birds recruited. The two observers positioned themselves at 10 m from the tripod at vantage points before launching the soundtrack with an MP4 player (NW-A45 Sony). Before launching any test, we made sure that no bird was already in the vicinity nor uttering mobbing calls in a distance. If a bird was detected, we waited only it left the area (~ 10 m around the loudspeaker). We observed the area with binoculars and all birds either calling and/or approaching from the beginning of the test to 15 sec after the end of the soundtrack. One bird was considered as approaching if it came in the 10 m radius around the tripod (Dutour et al. 2017b). Only birds uttering specific and known mobbing calls (see Sup. File 4 for spectrograms) were noted as calling. If a bird displayed the complete sequence of mobbing behavior (i.e., simultaneously calling and approaching the loudspeaker), it was then considered as giving a mobbing response. The two observers agreed on the highest number of birds seen simultaneously by both experimenters.

### Statistical analyses

All statistical analyses were done with R studio (R v.4.1.1, R core team 2022).

Since social conditions for our study species differ between winter and spring and factors influencing rates of response presumably therefor differ, the analysis was done separately for each season. We used three count response variables: the number of responding birds of any species (“community level”), the number of responding coal tits, and the number of responding crested tits. Given the high densities of both species in the study area, we considered that the absence of responding birds is due to the absence of response (i.e., structural zero) rather than the absence of bird (i.e., sampling zero). We therefore used Hurdle mixed models which are more convenient than zero inflation models to handle an excess of zeros of count data in such a situation (Zuur et al. 2009, Feng 2021). More specifically, Hurdle models are two stage models using a Bernoulli probability mass function to treat the zero outcomes as the result of a first process driving the occurrence of response (in our case, the mobbing occurrence), and a left truncated probability mass function to treat the positive outcomes as the result of a second process driving the response intensity (in our case, the intensity of mobbing). For each count variable, we first constructed an initial full Hurdle model implemented in the package *glmmTMB* (v.1.1.2.3, Brooks et al. 2017), with the effect of the emitter species, the effect of the number of callers, and their interactive effect in both parts of the model (occurrence and intensity). Moreover, both the spot location and the soundtracks’ ID were introduced as random effects as an intercept in the model. All models were constructed with a quasi-Newton optimization method (‘BFGS’) to circumvent convergence failure. Nevertheless, the random effects were discarded from the model when analyzing the response of crested tits because of a general lower response precluding the correct estimation of the random effects. In order to control for potential overdispersion in our positive count data, we first selected between two alternative left truncated probability mass functions to handle positive counts, a truncated Poisson distribution and a truncated negative binomial one allowing the variance to increase more rapidly than the esperance (note that we tested both nbinom1 and nbinom2, the former having a linear parameterization and the second having a quadratic parameterization, Hardin & Hilbe 2007). For this purpose, both models were constructed and compared using Bayesian Information Criterion (BIC) and AIC. Since BIC is more sensitive to the sample size but less sensitive to the unobserved heterogeneity than AIC (Brewer et al 2016), we only reported BIC. For the community response, a truncated negative binomial distribution led to the lowest BIC and was therefore chosen. Indeed, the dispersion parameter θ (i.e., the inflation factor associated to the truncated negative binomial distribution: when θ → 0, the distribution is closer to a Gamma distribution, while when θ → ^+^∞, the distribution is closer to a Poisson distribution) was 1.16 for the community model in winter and 0.79 for the community model in spring. For the isolated response of coal tits and crested tits, the truncated Poisson distribution led to the lowest BIC and was therefore chosen to analyze these responses. The fit of the structure selected for the initial model was then checked by the inspection of its residuals using the package DHARMa (v.0.4.5, Hartig and Hartig 2017).

For each of the three response variables (at the level of community, crested tits and coal tits) and for each season, we then created four candidate models, each of them with all the explanatory terms of interest (number of callers and emitter species), but for which the interaction term was kept or not, in the occurrence part and the intensity part of the model. Weighted BIC (wBIC) was then computed for the four candidate models and used to assess and compare their relative support using evidence ratios (i.e., ratio of wBIC between two models, Anderson and Burnham 2002). Effects sizes of the differences between treatments were calculated with odds ratios (OR).

### Ethical note

We used a sample size that is higher than in other recent studies (commonly around 20-30 tests per treatment) to circumvent common problems of lack of power in animal behavior studies, and because presence/absence data usually require larger sample sizes (Jennions 2003). To limit the impact on birds’ welfare, we ran short playback tests (1 min-long). All birds returned to a foraging behavior in less than 5 minutes after our tests. No direct contact between birds and humans nor any concealment of the birds were needed in this experiment.

## RESULTS

### Mobbing responses in winter

Eleven different species were attracted to our soundtracks (Figure 1A), with a maximum diversity of six species at one test. The four main species were the goldcrest (*Regulus regulus*, present in 29.5% of our tests), the crested tit (present in 27.8% of our tests), the coal tit (26.7%) and the marsh tit (*Poecile palustris*, 16.3%). As indicated by the best supported model (lowest BIC and an evidence ratio of 8.3, Table 1A), mobbing occurrence (the probability that at least one bird responded the playback), irrespective of the species (i.e., at the community level, Figure 2A) was affected by an additive effect of both the number of callers in the playback and the caller species (Table 2A). Indeed, birds mobbed more often the coal tit soundtracks compared to the crested tit soundtracks, and more to soundtracks with three birds rather than only one bird calling (1CO: 64%, 3CO: 77%, 1CR: 30%, 3CR: 59%). This additive effect was also detected when looking at the mobbing intensity (i.e., the number of mobbing birds when mobbing occurs, Figure 2B, Table 2A). Indeed, the largest mobs were initiated by playbacks with three coal tits (4.01 ± 3.17 birds, mean ± standard deviation, with a maximum of 15 birds) while the smaller mobs were initiated by playbacks with one crested tit (1.90 ± 1.21 birds).

**Figure 1.**
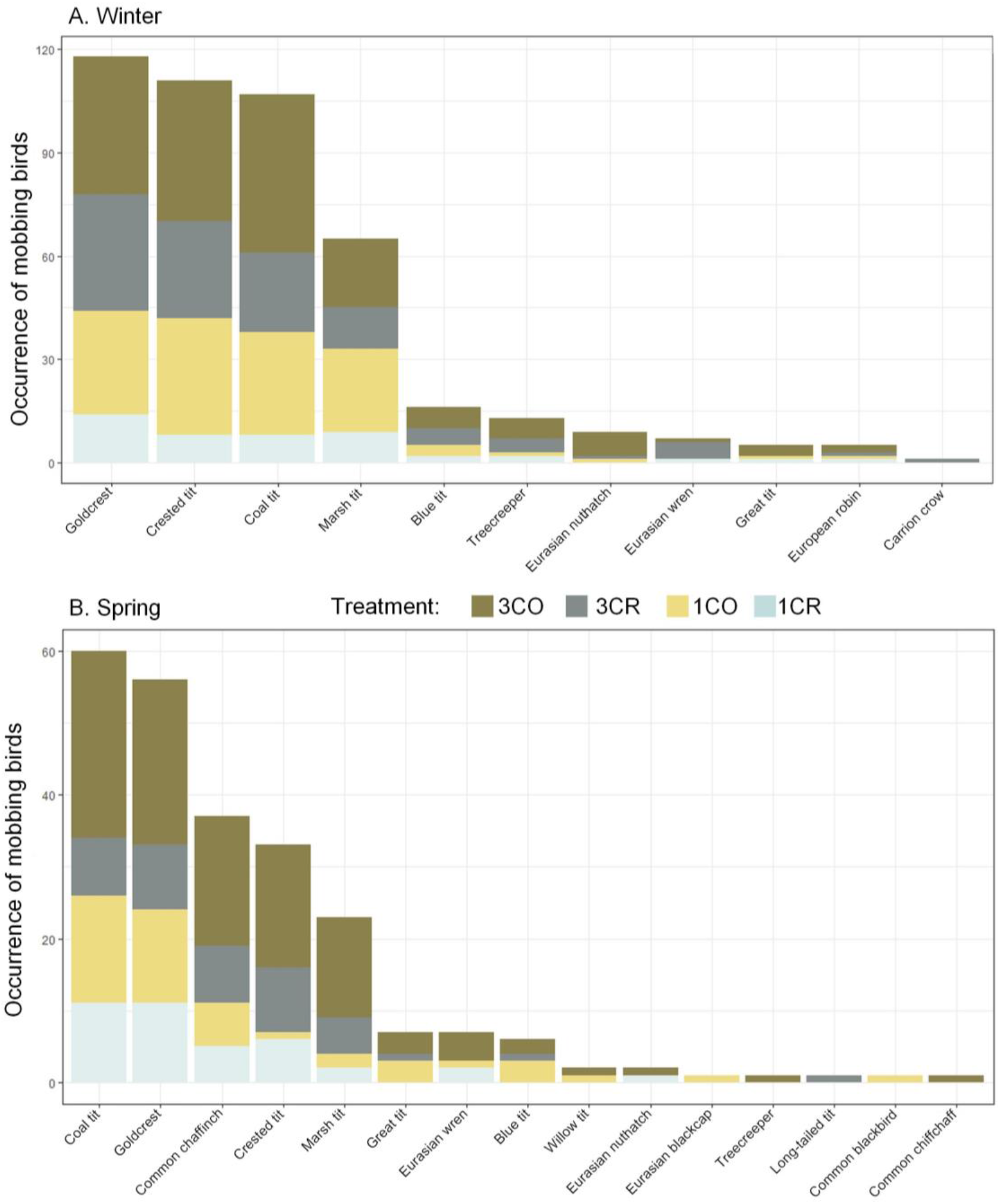
Number of spots (100 per acoustic treatment) in which at least one bird of each species mobbed (i.e., approached and called), when facing each of our four acoustic treatments (3CO: three coal tits, 1CO: one coal tit, 3CR: three crested tits, 1CR: one crested tit). Responses to each of the four treatments are stacked in sequence on each bar so that the entire bar represents the sum of all responses by a given species across treatments. Species taxonomy: blue tit = Cyanistes caeruleus, carrion crow = Corvus corone, crested tit = Lophophanes cristatus, coal tit = Periparus ater, common blackbird = Turdus merula, common chaffinch = Fringilla coelebs, common chiffchaff = Phylloscopus collybita, Eurasian nuthatch = Sitta europaea, Eurasian wren = Troglodytes troglodytes, Eurasian blackcap = Sylvia atricapilla, European robin = Erithacus rubecula, goldcrest = Regulus regulus, great tit = Parus major, long-tailed tit = Aegithalos caudatus, marsh tit = Poecile palustris, treecreeper = Certhia familiaris, willow tit = Poecile montanus.

**Table 1.**
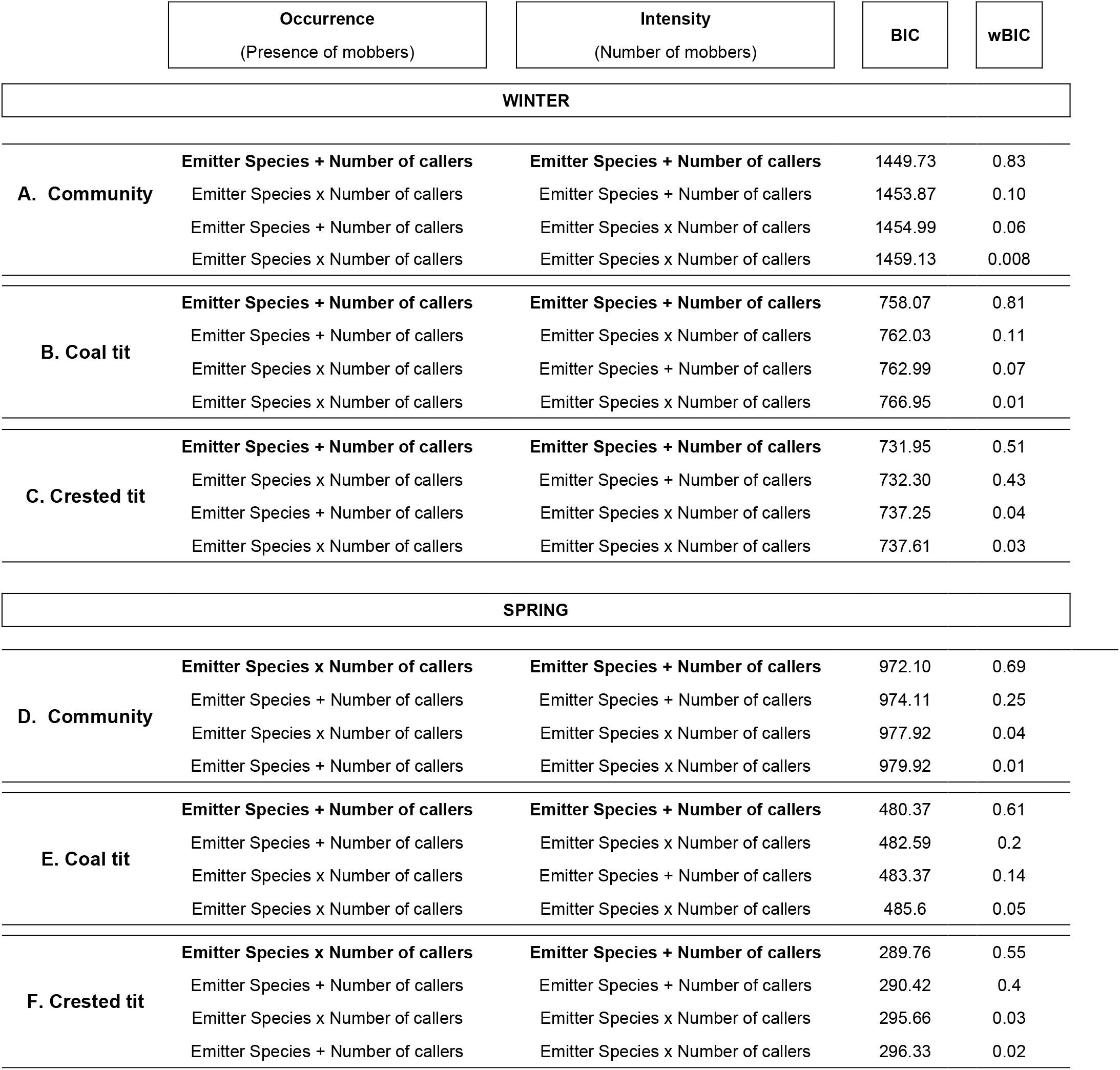
Hurdle models selection tables. For each response variable (number of responding birds at the community level, number of responding coal tits, number of responding crested tits), we first constructed a full Hurdle model with the effect of the emitter species, the effect of the number of callers as well as their interactive effect in both parts of the model (see Material and Methods for details). We compare this full model to models without the interaction in both the occurrence part and the intensity part of the model. We provide the Bayesian Information Criterion (BIC) and the weighted BIC (wBIC) to represent the relative support of each model. With wBIC we can calculate evidence ratio between two models (e.g., the first model is 0.83 / 0.10 ≈ 8.3 times more supported than the second model).

|  | Occurrence<br>(Presence of mobbers) | Intensity<br>(Number of mobbers) | BIC | wBIC |
| --- | --- | --- | --- | --- |
| **WINTER** |  |  |  |  |
| **A. Community** | **Emitter Species + Number of callers** | **Emitter Species + Number of callers** | 1449.73 | 0.83 |
|  | Emitter Species x Number of callers | Emitter Species + Number of callers | 1453.87 | 0.10 |
|  | Emitter Species + Number of callers | Emitter Species x Number of callers | 1454.99 | 0.06 |
|  | Emitter Species x Number of callers | Emitter Species x Number of callers | 1459.13 | 0.008 |
| **B. Coal tit** | **Emitter Species + Number of callers** | **Emitter Species + Number of callers** | 758.07 | 0.81 |
|  | Emitter Species + Number of callers | Emitter Species x Number of callers | 762.03 | 0.11 |
|  | Emitter Species x Number of callers | Emitter Species + Number of callers | 762.99 | 0.07 |
|  | Emitter Species x Number of callers | Emitter Species x Number of callers | 766.95 | 0.01 |
| **C. Crested tit** | **Emitter Species + Number of callers** | **Emitter Species + Number of callers** | 731.95 | 0.51 |
|  | Emitter Species x Number of callers | Emitter Species + Number of callers | 732.30 | 0.43 |
|  | Emitter Species + Number of callers | Emitter Species x Number of callers | 737.25 | 0.04 |
|  | Emitter Species x Number of callers | Emitter Species x Number of callers | 737.61 | 0.03 |
| **SPRING** |  |  |  |  |
| **D. Community** | **Emitter Species x Number of callers** | **Emitter Species + Number of callers** | 972.10 | 0.69 |
|  | Emitter Species + Number of callers | Emitter Species + Number of callers | 974.11 | 0.25 |
|  | Emitter Species x Number of callers | Emitter Species x Number of callers | 977.92 | 0.04 |
|  | Emitter Species + Number of callers | Emitter Species x Number of callers | 979.92 | 0.01 |
| **E. Coal tit** | **Emitter Species + Number of callers** | **Emitter Species + Number of callers** | 480.37 | 0.61 |
|  | Emitter Species + Number of callers | Emitter Species x Number of callers | 482.59 | 0.2 |
|  | Emitter Species x Number of callers | Emitter Species + Number of callers | 483.37 | 0.14 |
|  | Emitter Species x Number of callers | Emitter Species x Number of callers | 485.6 | 0.05 |
| **F. Crested tit** | **Emitter Species x Number of callers** | **Emitter Species + Number of callers** | 289.76 | 0.55 |
|  | Emitter Species + Number of callers | Emitter Species + Number of callers | 290.42 | 0.4 |
|  | Emitter Species x Number of callers | Emitter Species x Number of callers | 295.66 | 0.03 |
|  | Emitter Species + Number of callers | Emitter Species x Number of callers | 296.33 | 0.02 |

**Figure 2.**
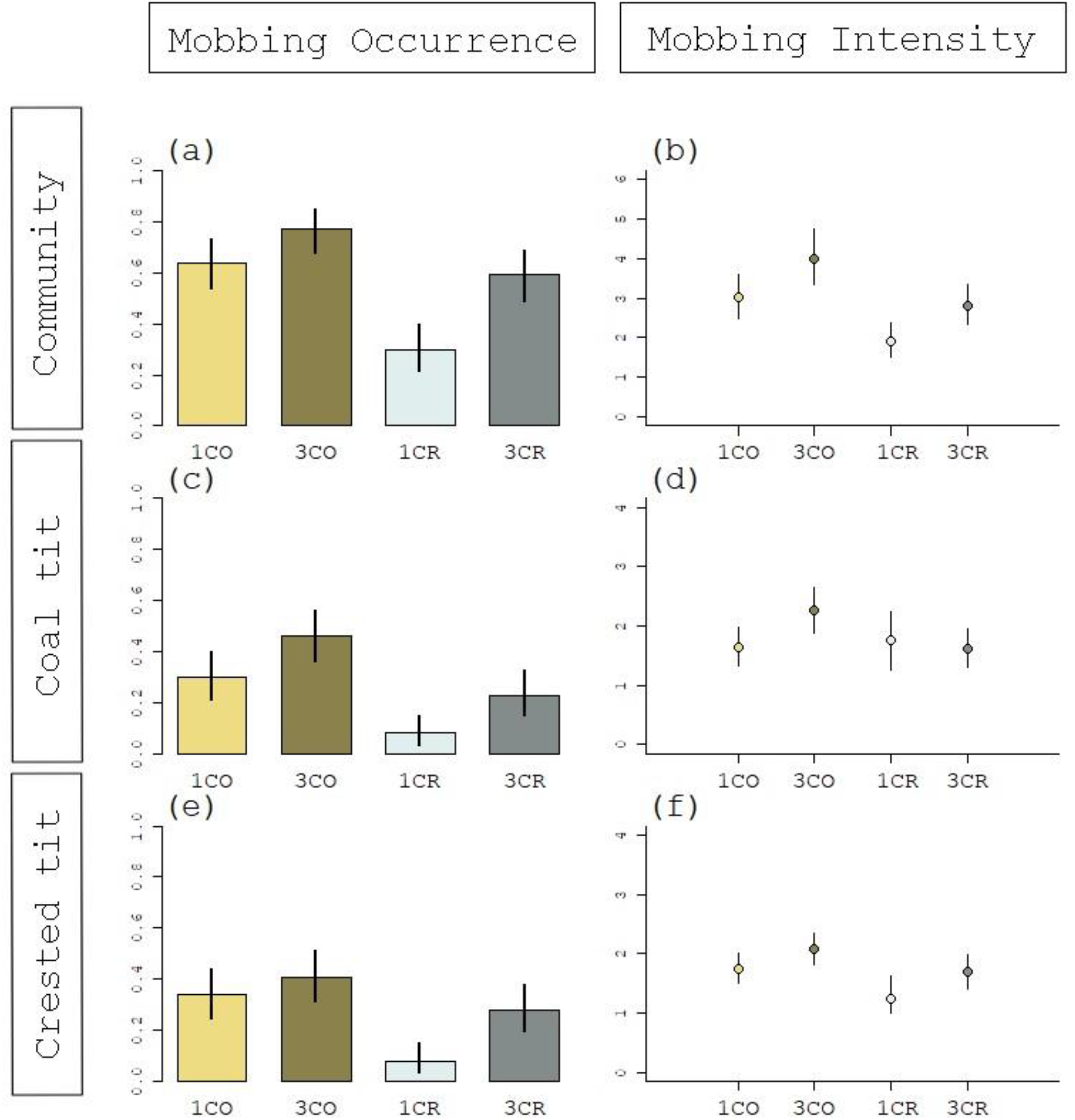
Mobbing response of the bird community tested in winter to our four different mobbing soundtracks (1CO: one coal tit, 3CO: three coal tits, 1CR: one crested tit, 3CR: three crested tits). Error bars are 95% confidence intervals. Graphs on the left represent mobbing occurrence: the proportion of spots in which at least one bird mobbed (i.e., approach and called, N = 100 per treatment). Graphs on the right represent mobbing intensity: the number of birds that responded when there was a mobbing response (sample sizes are the proportions of the graphs on the left). The upper graphs are responses of the general bird community, middle graphs are responses from coal tits, and lower graphs are responses from crested tits.

**Table 2.** Outputs of the models selected in bold in Table 1. Each Hurdle model is a two-stage model, the first one examining the effect of explanatory terms on the occurrence of response (mobbing occurrence) and the second one examining the effect of explanatory terms on the positive counts (mobbing intensity), see Material and Methods for details. We provide the estimates with their standard error (the intercept is the 1CO treatment), the z value and the associated p-value. The detailed outputs of the three other models in Table 1 are added as Sup. File 5.

| **WINTER** |  |  |  |  |  |
| --- | --- | --- | --- | --- | --- |
| **A. Community** |  | **Estimate** | **SE** | **z** | **p** |
|  | **Occurrence** |  |  |  |  |
|  | (Intercept) | -0.94 | 0.17 | -5.51 | < 0.0001 |
|  | Emitter Species | 1.19 | 0.23 | 5.23 | < 0.0001 |
|  | Number of Callers | 0.69 | 0.16 | 4.34 | < 0.0001 |
|  | **Intensity** |  |  |  |  |
|  | (Intercept) | 0.98 | 0.11 | 8.81 | < 0.0001 |
|  | Emitter Species | -0.58 | 0.16 | -3.67 | 0.0002 |
|  | Number of Callers | -0.33 | 0.11 | -3.06 | 0.0002 |
| **B. Coal tit** |  |  |  |  |  |
|  | **Occurrence** |  |  |  |  |
|  | (Intercept) | 0.53 | 0.16 | 3.32 | 0.001 |
|  | Emitter Species | 1.28 | 0.26 | 4.96 | < 0.0001 |
|  | Number of Callers | 0.63 | 0.18 | 3.57 | 0.0003 |
|  | **Intensity** |  |  |  |  |
|  | (Intercept) | 0.40 | 0.11 | 3.47 | 0.001 |
|  | Emitter Species | -0.44 | 0.23 | -1.87 | 0.06 |
|  | Number of Callers | -0.31 | 0.15 | -2.03 | 0.04 |
| **C. Crested tit** |  |  |  |  |  |
|  | **Occurrence** |  |  |  |  |
|  | (Intercept) | 0.53 | 0.15 | 3.55 | 0.0004 |
|  | Emitter Species | 1.03 | 0.24 | 4.33 | < 0.0001 |
|  | Number of Callers | 0.51 | 0.17 | 3.06 | 0.002 |
|  | **Intensity** |  |  |  |  |
|  | (Intercept) | 0.35 | 0.12 | 3.04 | 0.002 |
|  | Emitter Species | -0.48 | 0.24 | -2.01 | 0.04 |
|  | Number of Callers | -0.28 | 0.15 | -1.80 | 0.07 |

| SPRING |  |  |  |  |  |
| --- | --- | --- | --- | --- | --- |
| D. Community |  | Estimate | SE | z | p |
|  | **Occurrence** |  |  |  |  |
|  | (Intercept) | 0.05 | 0.16 | 0.29 | 0.77 |
|  | Emitter Species | 0.59 | 0.22 | 2.76 | 0.006 |
|  | Number of Callers | 0.97 | 0.22 | 4.43 | < 0.0001 |
|  | Emitter Species: Number of Callers | -0.90 | 0.30 | -2.97 | 0.003 |
|  | **Intensity** |  |  |  |  |
|  | (Intercept) | -0.08 | 0.44 | -0.19 | 0.85 |
|  | Emitter Species | -2.30 | 2.76 | -0.83 | 0.41 |
|  | Number of Callers | -0.55 | 0.36 | -1.54 | 0.12 |
| E. Coal tit |  |  |  |  |  |
|  | **Occurrence** |  |  |  |  |
|  | (Intercept) | 1.36 | 0.18 | 7.74 | < 0.0001 |
|  | Emitter Species | 0.90 | 0.30 | 3.02 | 0.003 |
|  | Number of Callers | 0.23 | 0.20 | 1.13 | 0.26 |
|  | **Intensity** |  |  |  |  |
|  | (Intercept) | -0.40 | 0.36 | -1.11 | 0.27 |
|  | Emitter Species | -0.97 | 0.63 | -1.53 | 0.13 |
|  | Number of Callers | -0.13 | 0.31 | -0.41 | 0.69 |
| F. Crested tit |  |  |  |  |  |
|  | **Occurrence** |  |  |  |  |
|  | (Intercept) | 3.09 | 0.52 | 5.95 | < 0.0001 |
|  | Emitter Species | -0.56 | 0.59 | -0.95 | 0.34 |
|  | Number of Callers | 2.13 | 0.74 | 2.90 | 0.004 |
|  | Emitter Species: Number of Callers | -1.82 | 0.83 | -2.19 | 0.03 |
|  | **Intensity** |  |  |  |  |
|  | (Intercept) | -0.06 | 0.48 | -0.12 | 0.90 |
|  | Emitter Species | -1.30 | 0.88 | -1.048 | 0.14 |
|  | Number of Callers | 0.51 | 0.61 | 0.84 | 0.40 |

When focusing on the occurrence of response of coal tits or the one of the crested tits, the best supported model comprised an additive effect of the number of callers and the emitter species (Table 1B and 1C, Table 2B and 2C), resulting in a lower response toward singletons of crested tits (8% of points attracted coal tits or crested tits), intermediate scores toward trios of crested tits and singletons of coal tits, and the highest occurrence of response toward soundtracks with three coal tits (46% triggered a response from coal tits and 41% triggered a response from crested tits, Figure 2C and 2E). However, for the crested tit, the model with an interaction between number of callers and emitter species was also well supported (evidence ratio of 0.51/0.43 = 1.19, Table 1C). Indeed, the difference between 1CR and 3CR was higher (OR: 4.74, 95%CI: [1.92; 10.40]) than the difference between 1CO and 3CO (OR: 1.35, 95%CI: [0.76; 2.40]). Regarding mobbing intensity (Figure 2D and 2F), for both the coal tits’ and crested tits’ response, the additive effect of number of callers and emitter species was less stringent than for the occurrence of mobbing (the effect of emitter species for the coal tit, and the effect of number of callers for the crested tit did not reach statistical significance when reporting the estimates, Table 2B and 2C).

### Mobbing responses in spring

In spring, we detected a lower mobbing propensity: 58% of our tests did not trigger any mobbing behavior, while this proportion was of 42.5% in winter. Fifteen different species were attracted to our soundtracks (Figure 1B), with a maximum diversity of four species at one test. The four most common species that responded were the coal tit (present in 15% of our tests), the goldcrest (present in 14% of our tests), the common chaffinch (*Fringilla coelebs*, 9.25%), and the crested tit (8%).

Regarding mobbing occurrence at the community level (Figure 3A), the model with the lowest BIC was the one including an interaction between emitter species and number of callers in the playbacks (Table 1D, Table 2D). Indeed, the effect sizes depict a higher response towards the 3CO treatment than towards any of the three other playbacks (e.g., 3CO vs 3CR: 3.30, 95%CI: [1.85; 5.89]), while the three other playbacks triggered a similar response (e.g., 3CR vs 1CO: 1.14, 95%CI: [0.64; 2.05]). Note however that this interaction is not strongly supported since the model including only the additive effects of number of callers and emitter species gave a similar BIC (evidence ratio of 0.69/0.25 = 2.76, Table 1D). When focusing on the intensity of response (Figure 3B), we detected no difference in the number of birds recruited to the four different playbacks (Table 2D). The number of birds in the mob never exceeded seven birds.

**Figure 3.**
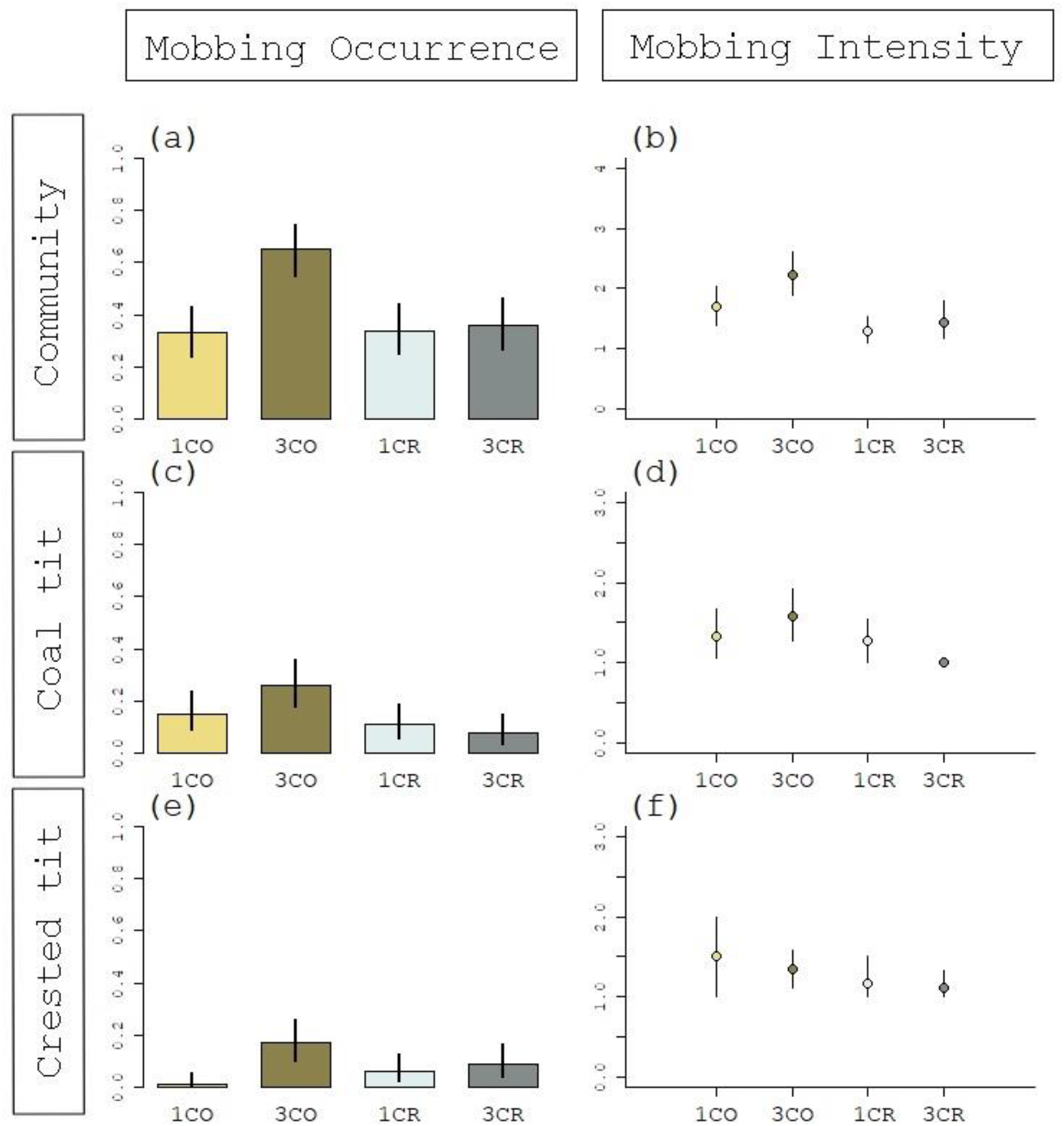
Mobbing response of the bird community tested in a replication of the first experiment, during the reproductive season (spring). Birds’ responses are recorded when facing four different mobbing soundtracks (1CO: one coal tit, 3CO: three coal tits, 1CR: one crested tit, 3CR: three crested tits). Error bars are 95% confidence intervals. Graphs on the left represent mobbing occurrence: the proportion of spots in which at least one bird mobbed (i.e., approach and called, N = 100 per treatment). Graphs on the right represent mobbing intensity: the number of birds that responded when there was a mobbing response (sample sizes are the proportions of the graphs on the left). The upper graphs are responses of the general bird community, middle graphs are responses from coal tits, and lower graphs are responses from crested tits.

The best supported model regarding the presence of at least one coal tit included the emitter species of the playback, but no effect of the number of callers (Figure 3C, Table 1E, Table 2E). For the crested tit’s occurrence, we recorded an interaction between the emitter species and the number of callers in the playbacks (Figure 3E, Table 1F, Table 2F). Indeed, our playbacks attracted more often crested tits when there were three coal tits in the playbacks compared to any of the three other types of playbacks. For both species, the number of birds recruited when mobbing occurred did not differ between the four types of playbacks (Figure 3D and 3F, Table 2E and 2F).

## DISCUSSION

In winter, coal tits’ soundtracks triggered more mobbing response from conspecifics and heterospecifics than crested tits’ soundtracks; and soundtracks with three callers triggered more mobbing response from the bird community than soundtracks with only one caller. However, when replicating the experiment in spring, we found a lower general response but also differences between playbacks, with increased responses only toward the 3 coal tits’ playbacks. This interaction between context and acoustic cues demonstrates the flexible nature of heterospecific communication.

### In winter, both the number of caller and emitter species influence mobbing responses

Birds often modulate their mobbing responses depending on the threat they perceive. For example, different predators are mobbed with different levels of intensity (Curio et al. 1983, Templeton et al. 2005). Individuals can also change their mobbing response depending on the distance of the threat, the movement of the predator, or other cues surrounding the predator (Book & Freeberg 2015, Carlson et al. 2017). In this study, we recorded a higher mobbing response towards soundtracks with three individuals than towards soundtracks with only one individual calling. This result is congruent with the hypothesis that birds use acoustic cues to gain information on the threat. Indeed, a larger number of birds may indicate a more significant predator, as larger mobs are produced in front of more important predators (Dutour et al. 2017b, Sandoval & Wilson 2012). In addition, joining a group instead of a lone caller increases the dilution effect, hence reducing risk for the newcomer (Sridhar et al. 2009). Alternatively, the increased response to the playbacks with more birds may be unrelated to an assessment of risk by birds, but rather be a simple mechanical threshold reached when the call is more salient to receivers (by reaching a specific threshold and/or being easier to detect). One solution to test the risk assessment hypothesis could be to create a similar experiment but based only on visual cues. The idea would be to test the mobbing response of birds in front of a predator model accompanied with either one or three models of conspecifics. This kind of experiment should be done in large aviaries for which we can control the visual cues the birds receive. If the focal bird approach and mob more a when a group is already present, then the risk hypothesis would be more supported.

The mechanisms implicated in the differentiation between playbacks of one and three callers can be various. In natural settings, birds can consider the number of spatially different acoustic sources (Bradbury and Vehrencamp 2011). In our study, we launched the soundtracks with only one loudspeaker whatever the treatment to suppress this effect. Therefore, in our tests, the acoustic criteria that remain available are the duty cycle (i.e., the proportion of the calling sequence when the signal is present), and the count of calling individual through individual signatures. Our experiment does not add any insights on which criteria was used by birds. Based on the current literature, the duty cycle is probably one major coding strategy for increased risk in Parids (Landsborough et al. 2020, Salis et al. 2022), and Parids modify their response to unknown non-Parids calls with different duty cycles (Dutour et al. 2022). Yet, great tits can also recognize caller identity, as they increased their mobbing response toward soundtracks made with calls of several individuals compared to soundtracks with only one individual calling (Dutour et al. 2021). In this latter experiment, the duty cycles of the different treatments were strictly equal. This result was however not replicated when testing the response to different number of heterospecifics (chaffinches *Fringilla coelebs*, Dutour and Randler 2021). In our experiments, we believe that the overlapping of the calls in the three birds treatments avoid the risk of interpreting these treatments as only one bird calling intensely. Further experiments exploring the response of each species to conspecific and heterospecific calls with controlled duty cycle may enlighten whether individual recognition can also be used in heterospecific communication.

A mobbing response occurred more often when broadcasting coal tits’ mobbing calls compared to crested tits’ calls, but also more birds responded to it. Unexpectedly, even crested tits responded more to coal tits’ mobbing calls than to calls from their own species. Coal tits therefore appear to be listened to and heavily responded to, leading to larger (and possibly more efficient) mobs. This is in line with the hypothesis that species from the same community show different levels of reliability (Magrath et al. 2015). The notion of “a community informant” was developed for Parids in Carlson et al. (2020). They investigated whether the birds possessed a reliable way of encoding predator information, and if several heterospecifics relied on these calls. They showed that the great tit (*Parus major*) best fitted the definition of community informant. The coal tit approached the definition, with only one caveat: the dunnock (*Prunella modularis*) did not respond to it. As the authors suggested, the lack of response from one species does not mean that other species from the community do not respond to it (Carlson et al. 2020). Indeed, in our study, 14 species responded to coal tits’ soundtracks. In contrast, the crested tit did not meet any of the criteria set by Carlson and colleagues. Coal tits appear therefore to be one important species regarding predator information in the community, and this is congruent with their increased sensibility to predation by pygmy owls (*Glaucidium passerinum*) in winter (Suhonen et al. 1993).

### Replicating the experiment in spring: A lower general response

In winter, Parids living in temperate regions often flock with heterospecifics, sometimes leading to impressive mobs (up to 20 birds in the present experiment). In opposition, during the reproductive period (May-July), Parids nest and defend their territory with intensity (Hinde 1952). For this reason, we first explored the mobbing response of birds in winter, as this is the season in which interactions and cooperative mobbing with heterospecifics makes more sense. However, we replicated the experiment in spring to explore whether seasonal context of the experiment could impact our results. We did not test the same birds and cannot control the changes in environment and community between the first tests in winter and the replicate in spring. For these reasons, we did not statistically compare the two seasons, but will nonetheless discuss the differences found between the original experiment and the replication.

In spring, the number of birds mobbing to the four different types of playbacks was lower than in winter and did not differ between playback types. We here propose that in spring, when all birds defend their territory, the number of birds that can respond is restricted to the neighbors. Moreover, in spring, aggressiveness toward conspecifics is high and may therefore reduce the number of potential birds responding to mobbing calls. This aggressiveness may also explain why not so many birds responded to conspecific mobbing calls in spring (coal tits to coal tits and crested tits to crested tits).

Additionally, not only did fewer individuals respond in spring than in winter, but in spring, the proportion of locations with any response was lower than in winter. This difference must be taken with cautiousness, as the community and the density of the populations may vary with the seasons: a decrease in mobbing response may simply be related to fewer individuals in the territory. An order effect due to tests in winter being done before the tests in spring is unlikely given the absence of order effect in our experiment at a short time scale (see Sup. File. 2 For details). In addition, in spring, we were able to hear coal tits singing at the 100 spots studied. We are therefore confident that, in spring, each spot could have recorded one coal tit’s mobbing response. This suggests that at least for the coal tit, the response to conspecific and heterospecific mobbing calls decreases in spring. This result is consistent with Dutour et al. (2019) who detected in Parids a higher mobbing response toward heterospecific calls in winter compared to summer. The proximal reasons for such a decrease can be various. Increased territoriality and aggression in spring may very well limit cooperative communication, since the mobbing calls may resemble intra-specific aggression/territoriality calls, leading to a lower relevance for heterospecifics. Other factors such as decreased predator pressure in spring (Dutour et al. 2017b) could also result in a lower investment in mobbing in spring. The ratio cost/benefits in responding to distanced mobbing calls is therefore probably flexible through different times of the year. Given that most of these factors are intercorrelated, determining which one is responsible for the difference in mobbing is unfeasible in natural conditions.

### Replicating the experiment in spring: Almost no response to crested mobbing calls

In addition to a general lower mobbing response in spring, the differences between treatments were also impacted by the season. Indeed, while we selected similar models for the community, coal tits’, and crested tits’ response in winter (additive effect of number of caller and emitter species), we found support for different models in spring. A general tendency was detected, with only the playbacks with three coal tits triggering more response than the three other playbacks. This suggests that the crested tit is not considered as informative in spring, even when mobbing in groups, and unexpectedly, even to conspecifics. Several explanations can be proposed. First, a group of three crested tits in spring may be too rare to bear meaning, as they are in pairs and defending their nest. However, this hypothesis does not stand as this is also the case for the coal tit, but that the difference between one and three callers still stands in spring for this species. Alternatively, the contact with crested tits may be reduced in spring if crested tits densities are lower during this season, hence decreasing learning opportunities for heterospecifics. However, crested tits stay on the same territory throughout the year (Ekman 1979) making this hypothesis unlikely despite the fact that our experiments do not allow us to formally rule out it. We rather suggest that this lack of mobbing response may emerge from reduced reliability of the calls. To be efficient, an acoustic signal needs to be easily distinguishable from other signals (Bradbury and Vehrencamp 2011). The song and mobbing calls of the crested tits are extremely similar (Cramp and Perrins 1993, Hailman 1989). As crested tits produce both songs and mobbing calls in spring, we can hypothesize that the global vocal production of crested tits therefore becomes less reliable from an external individual, hence leading to a decreased response to such calls. In contrast, the coal tit appears to be reliable and responded to in both seasons. This result is consistent with Jiang et al. (2020) who also found that between seasons differences in playback responses did not affect the nuclear status of some particular species (in their case, David’s fulvetta *Alcippe davidí*). The difference between the response of the bird community to coal tits’ playbacks compared to crested tits’ playbacks may also be due to a higher aggressiveness from crested tits. Crested tits are known to be more aggressive during spring (Campbell 1958), and crested tits, larger than coal tits, have higher rank dominance status (Suhonen et al. 1993). We have, however, little data on whether the heterospecific aggressiveness is higher than coal tits’ aggressiveness since dominance status is not necessarily linked to increased aggressiveness (Wilson 1992). Finally, difference in nest predation may impact the reliability of the information produced, but to our knowledge, nest predators are similar between Parid species (Cramp and Perrins 1993).

To conclude, birds from a community respond differently to acoustic situations with varying emitter species and number of callers. The number of callers may be recognized either with caller identity and/or changes in duty cycles. Those acoustic cues are not responded to in the same way throughout the year, possibly because of changes in territoriality and reliance on heterospecific calls. These results emphasize the importance of seasons in studies investigating the complexity of heterospecific communication.

## Supporting information

All supplementary files cited in the text (also on Zenodo)

## Acknowledgments – Funding

This work was supported by the French Ministry of Research and Higher Education funding (to A.S. PhD grants 2019-2022). We are grateful to Coline Guignier and Lou Sayd for useful help with fieldwork. We thank the *Ligue de Protection des Oiseaux* (LPO AuRA) for the information on the presence of coal and crested tits in the territory. We are thankful to the two reviewers and the recommender of PCI for their useful comments on the manuscript.

## Ethics

All the authors follow the ASAB/ABS Guidelines for the Use of Animals in Research and with The European Code of Conduct for Research Integrity. This work did not require special permit. Conflict of interest: none.

## Data statement

Raw data, R code, and Supplementary files are all added in Zenodo. https://doi.org/10.5281/zenodo.7649854

