## Supplementary material for "Acoustic cues and season affect mobbing responses in a bird community": All supplementary files cited in the text (also on Zenodo)

**Supplementary File 1: Testing the independence of the playback tests.**

In the present study, we used 100 points spaced at a minimum of 100 meters. Such a distance is sufficient to degrade bird sounds in a forest (although not perfectly as different frequencies degrade differently, Morton 1975), but birds may still follow the experimenters from one test place to the following when they hear the next soundtrack. Consequently, experimenters may test the same bird several times and therefore obtain data with pseudoreplication. While we made sure to check that no bird was following us, we agree that some birds may pass through our observations. We therefore ran a short experiment to test whether coal and crested tits could be tested several times in our protocol. To this aim, we went back to the same points selected in our study in late October 2021, when the birds are in flocks. We did not recreate the same protocol in summer, with the hypothesis that movements of birds are even more constrained during the reproductive season than in winter due to their nest place and breeding activities.

At each point, a soundtrack was launched, and two experimenters observed the birds approaching and calling for ~ 1 min at a distance. Experimenter A then took the loudspeaker and went to the next point. Experimenter B stayed and chose one bird (either coal or crested tit) as a focal bird and followed it for 5 min (average time between two tests in our study). At the end of the 5 min, experimenter A launched a second soundtrack while experimenter B recorded the focal bird's response when hearing the soundtrack. This protocol was repeated 22 times (9 crested and 13 coal tits). Although experimenter B focused on only one bird, they often came in flocks. Consequently, a total of 33 coal tits, 21 crested tits, and 7 other heterospecifics were followed.

Out of these 22 tests, no bird moved farther than 50 m from the loudspeaker during the 5 min observations. Most birds stayed on nearby trees ~15 m from the loudspeaker, gradually decreased their call rate during the 5 min period when they were observed. They never approached the second loudspeaker when it was launched. On few occasions (4/22), birds made few calls but with a lower intensity than the first time. A similar experiment was done by Salis et al. 2022 with the same conclusions, but with the Great tit *Parus major*. It seems therefore improbable that a tested bird was tested twice on two consecutive tests. We can fairly assume that our tests were independent.

**Supplementary File 2: Testing a potential order effect**

In this experiment, we carried out at each season four tests at each chosen point (100 points), in a short time span (2 weeks). We alternated our treatments at each consecutive points, hence creating four groups of 25 points with different order in their treatment (i.e., the first group first received 1CO, then 3CO, then 1CR and lastly 3CR, while the second one received 3CR -> 1CO -> 3CO -> 1CR, etc). In this additional analysis, we test whether order of playbacks had an impact in our results. Two kinds of order could impact our results: 1/whether one treatment, more than the others, impact the following treatment received, and 2/ whether the repeated playback stimulation could decrease birds' response by habituation.

We therefore ran two models per season with the total number of birds mobbing at the end of the test as the response variable. For the first model, we added treatment, order the treatment (i.e., whether birds first heard the 1CO calls or one of the other three treatments) and their interaction was added as explicative variables. For the second model, we added treatment, order in the timeline (i.e., whether the test was done on the first occasion we went of the field up to the fourth time), and their interaction. We used a generalized linear model with a Poisson distribution. However, for the two models in Winter, we detected an overdispersion and therefore used a negative binomial distribution instead. We estimated the importance of each term with function *Anova* and calculated pairwise comparisons with *emmeans*.

In winter, the only significant term was the treatment in both models: neither the effect of order of playback presentation nor the effect of repeated playback presentations impacted the birds' behavior. In Spring, for both models we detected an interaction between order and treatment. However, when investigating which groups were different inside each treatment, we did not find any statistically different pairwise comparisons, even without any correction for multiple comparisons. We can therefore conclude that the occurrence of birds in our tests was not impacted neither the repeated presentations nor the order in which playbacks were presented.

**Table 1.** Models with the number of presentations of playbacks or order of the different treatments on the total number of birds mobbing in response to playbacks. GLM with Poisson distribution are used in Spring, and with a negative binomial distribution in Winter.

| | SPRING | | | WINTER | | |
|---|---|---|---|---|---|---|
| | $X^2$ | Df | P value | $X^2$ | Df | P value |
| **Repeated playbacks** | | | | | | |
| Treatment | 76.23 | 3 | < 0.001 | 76.34 | 3 | < 0.001 |
| Repetition | 4.51 | 3 | 0.21 | 4.85 | 3 | 0.18 |
| Treatment*Repetition | 10.94 | 9 | 0.28 | 17.51 | 9 | 0.04 |
| **Order of playbacks** | | | | | | |
| Treatment | 76.23 | 3 | < 0.001 | 74.66 | 3 | < 0.001 |
| Order | 0.78 | 3 | 0.85 | 0.96 | 3 | 0.81 |
| Treatment*Order | 14.67 | 9 | 0.10 | 21.39 | 9 | 0.01 |

**Figure 1.** Number of birds mobbing depending on the different treatment and (A) the order of presentation of the playbacks and (B) the number of presentation of playbacks. Mean and 95% confidence intervals are given.

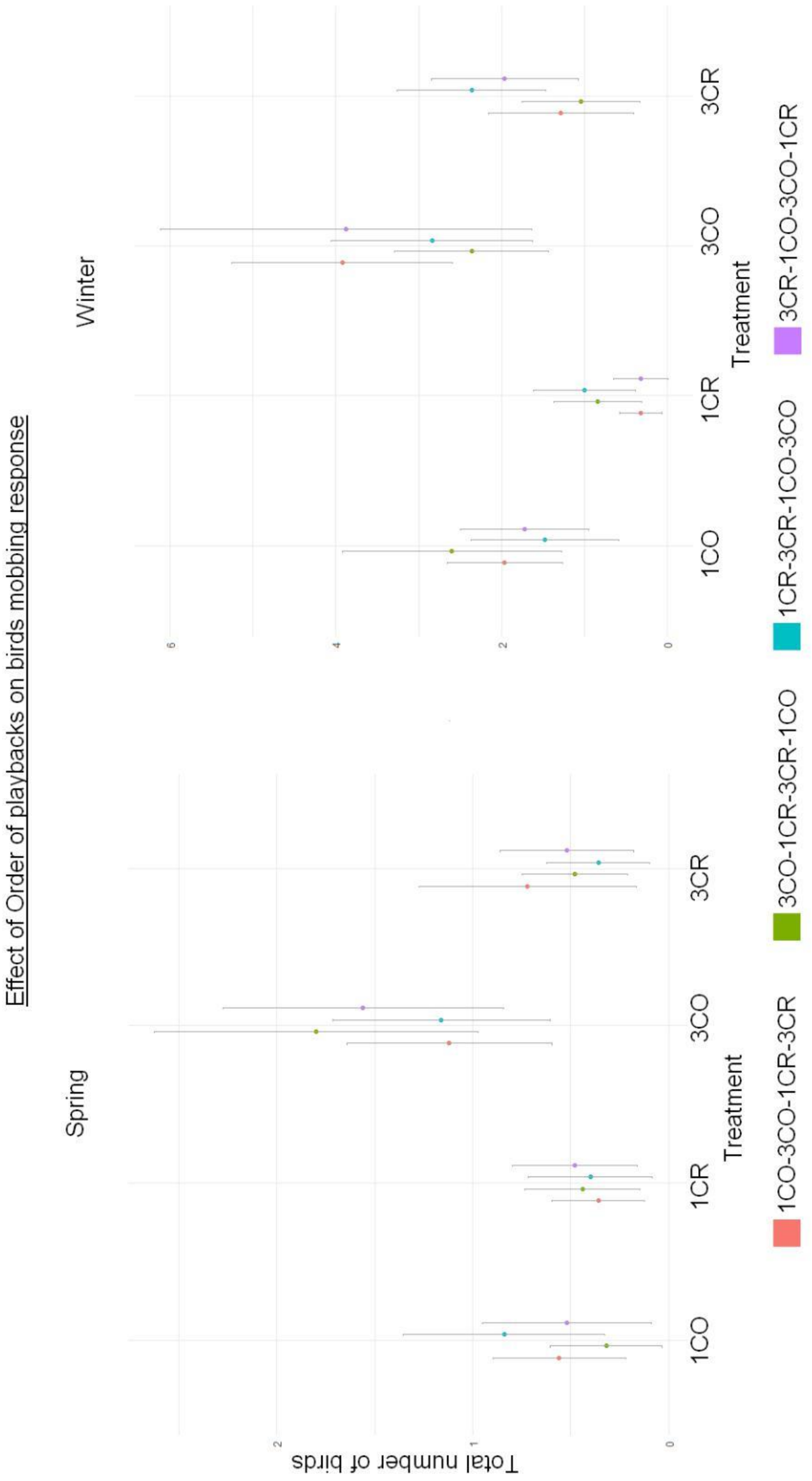

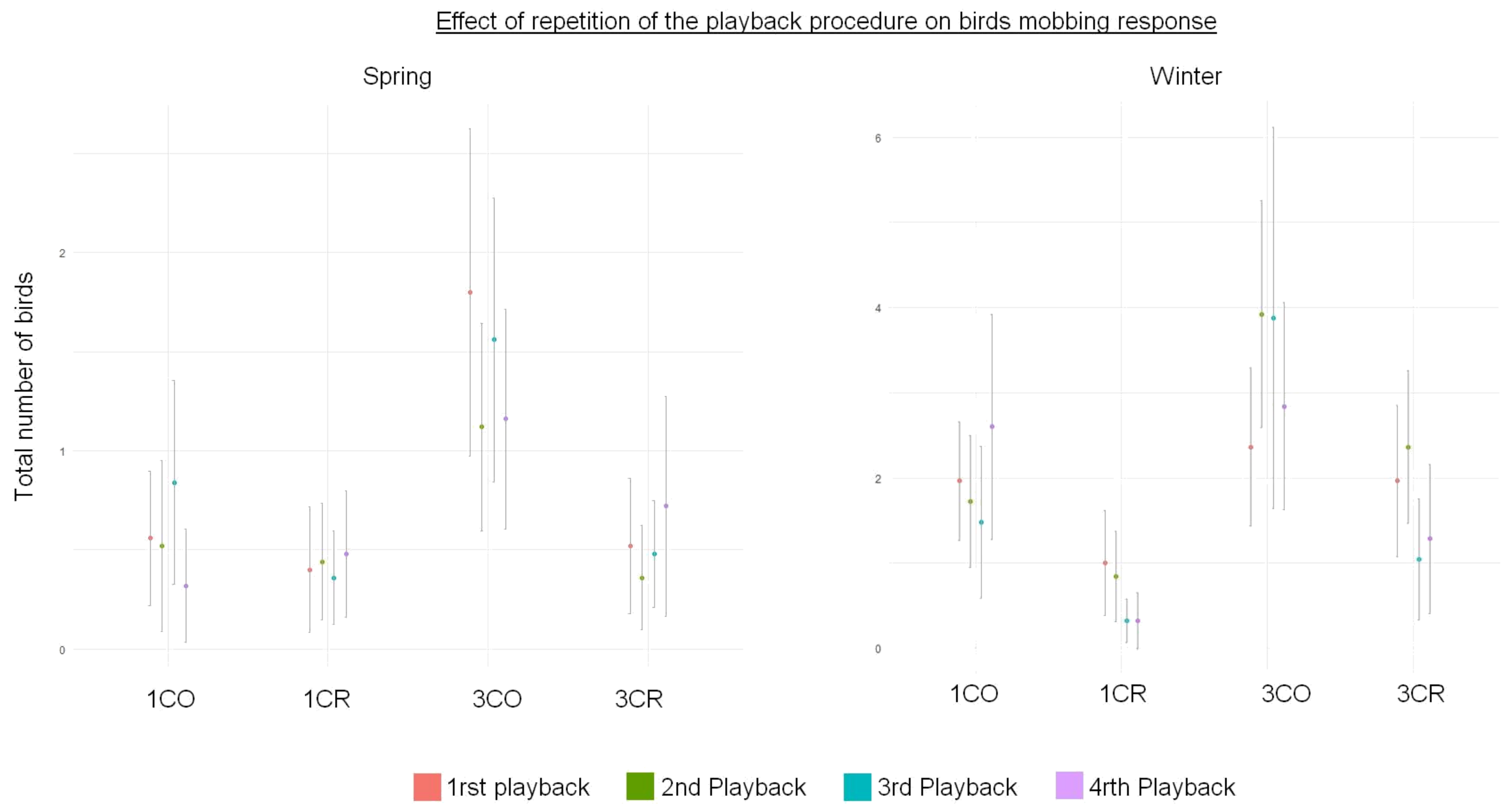
Effect of repetition of the playback procedure on birds mobbing response
Spring
Winter
Total number of birds
0
1
2
4
6
1CO
1CR
3CO
3CR
1rst playback
2nd Playback
3rd Playback
4rth Playback

**Supplementary File 3.** Additional information about playbacks of coal and crested tit used in the study.

We simulated mobbing choruses of three birds by superimposing three independent recordings. However, for the crested tits, we often possessed recordings with already two birds calling simultaneously. We therefore only added one more individual for 3CR_A, 3CR_C, 3CR_D, and 3CR_E.

The coal tit produces long notes that are not specifically clustered into calls; while the crested tit produced short notes clustered into defined calls. Hence, we report here the number of notes in each playback, which is therefore stronger for the crested tit. We controlled our playbacks by duty cycle, i.e., the amount of signal in the recordings. The duty cycle was calculated with Avisoft SASLab by removing the silent sections from the files, hence giving a duration in seconds.

<table>
<tr><th>Final File</th><th>Original File ID</th><th>Number of birds</th><th>Nb of notes</th><th>Final duty cycle (seconds)</th><th>Duty cycle : Mean ± standard deviation for each treatment</th></tr>
<tr><td rowspan="3">3CO_A</td><td>1</td><td>1</td><td>41</td><td rowspan="3">11.5</td><td rowspan="15">9.78 ± 1.18</td></tr>
<tr><td>2</td><td>1</td><td>42</td></tr>
<tr><td>3</td><td>1</td><td>39</td></tr>
<tr><td rowspan="3">3CO_B</td><td>1</td><td>1</td><td>45</td><td rowspan="3">9</td></tr>
<tr><td>2</td><td>1</td><td>64</td></tr>
<tr><td>3</td><td>1</td><td>33</td></tr>
<tr><td rowspan="3">3CO_C</td><td>1</td><td>1</td><td>45</td><td rowspan="3">10</td></tr>
<tr><td>2</td><td>1</td><td>49</td></tr>
<tr><td>3</td><td>1</td><td>33</td></tr>
<tr><td rowspan="3">3CO_D</td><td>1</td><td>1</td><td>40</td><td rowspan="3">8.4</td></tr>
<tr><td>2</td><td>1</td><td>53</td></tr>
<tr><td>3</td><td>1</td><td>31</td></tr>
<tr><td rowspan="3">3CO_E</td><td>1</td><td>1</td><td>57</td><td rowspan="3">10</td></tr>
<tr><td>2</td><td>1</td><td>50</td></tr>
<tr><td>3</td><td>1</td><td>52</td></tr>
</table>

| | | | | | |
|---|---|---|---|---|---|
| **1CO_A** | 1 | 1 | 42 | 4.4 | 6.64 ± 2.25 |
| **1CO_B** | 1 | 1 | 46 | 5.7 | |
| **1CO_C** | 1 | 1 | 46 | 9 | |
| **1CO_D** | 1 | 1 | 38 | 4.5 | |
| **1CO_E** | 1 | 1 | 40 | 10.1 | |
| **3CR_A** | 1 | 1 | 135 | 11.6 | 8.92 ± 2.64 |
| | 2 | 2 | 141 | | |
| **3CR_B** | 1 | 1 | 113 | 6.5 | |
| | 2 | 1 | 109 | | |
| | 3 | 1 | 142 | | |
| **3CR_C** | 1 | 2 | 275 | 11 | |
| | 2 | 1 | 172 | | |
| **3CR_D** | 1 | 1 | 148 | 7.5 | |
| | 2 | 2 | 247 | | |
| **3CR_E** | 1 | 2 | 242 | 8 | |
| | 2 | 1 | 165 | | |
| **1CR_A** | 1 | 1 | 168 | 8.5 | 6.98 ± 1.51 |
| **1CR_B** | 1 | 1 | 172 | 8.4 | |
| **1CR_C** | 1 | 1 | 129 | 5.6 | |
| **1CR_D** | 1 | 1 | 100 | 5.3 | |
| **1CR_E** | 1 | 1 | 159 | 7.1 | |

**Supplementary File 4: Spectrograms of the mobbing calls of the coal tit and the crested tit, and mobbing calls of the three most common heterospecifics that joined mobbing choruses.**

Created with Avisoft SASLab, FFTLength 512, Bandwidth 324 Hz, Resolution 86 Hz.

*Crested tit (Lophophanes cristatus)*

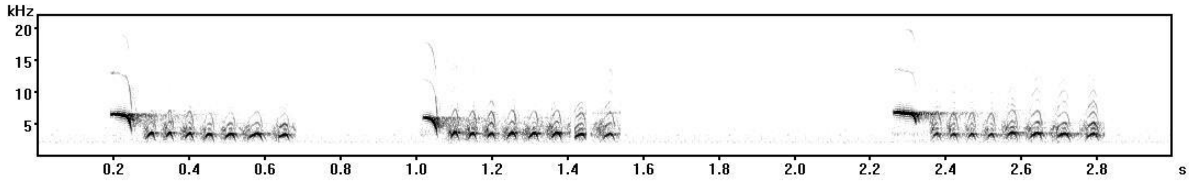


*Coal tit (Periparus ater)*

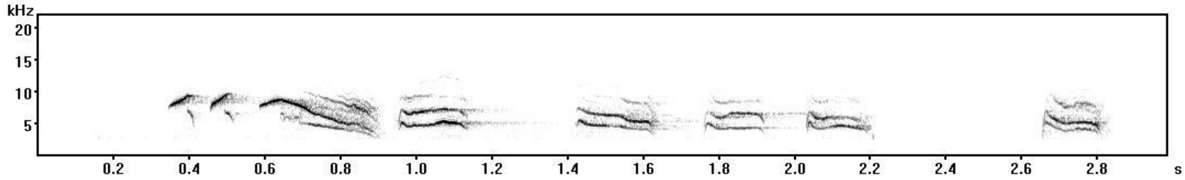


*Marsh tit (Poecile palustris)*

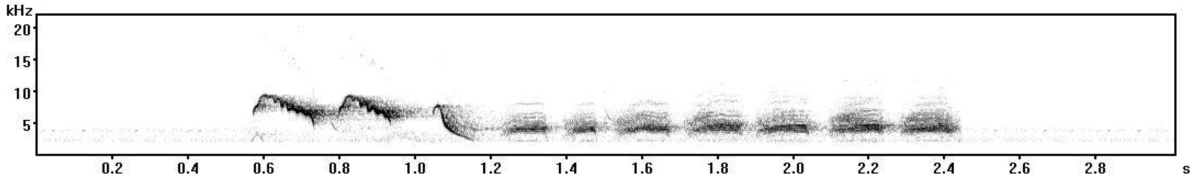


*Common chaffinch (Fringilla coelebs)*

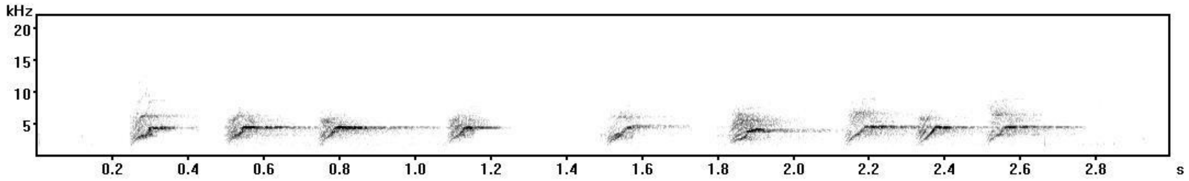


*Goldcrest (Regulus regulus)*

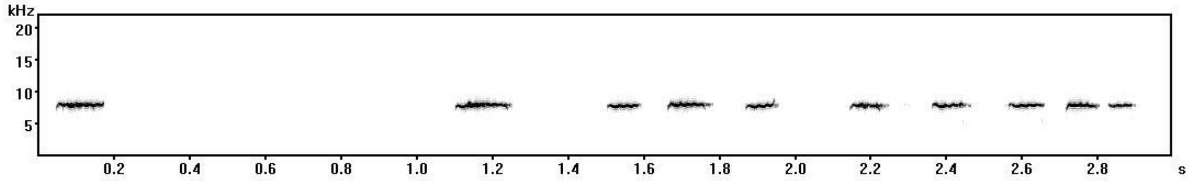

**Supplementary File 5: Model outputs**

Outputs of three models for each season and each variable that were not selected for presentation in the results (higher BIC than the selected model). The ID of each model (Hurdle #) are a reference to the names of the models that can be found in the R code provided with this manuscript. We provide the models for the community response in winter (A), in spring (B); for the coal tits' response in winter (C) and in spring (D) and for the crested tits' response in winter (E) and in spring (F). For a complete description of how we built our Hurdle models, please refer to the Material & Methods section of the main text.

### COMMUNITY

#### A. Winter

##### Hurdle 1

Occurrence

| | Estimate | SE | z | p |
|---|---|---|---|---|
| (Intercept) | -0,92 | 0,17 | -5,42 | <0,0001 |
| Emitter Species | 1,17 | 0,22 | 5,16 | <0,0001 |
| Number of Callers | 0,46 | 0,23 | 2,03 | 0,04 |
| Emitter Species * Number of Callers | 0,43 | 0,31 | 1,36 | 0,17 |

Intensity

| | Estimate | SE | z | p |
|---|---|---|---|---|
| (Intercept) | 1,00 | 0,11 | 8,92 | <0,0001 |
| Emitter Species | -0,63 | 0,17 | -3,71 | <0,0001 |
| Number of Callers | -0,27 | 0,13 | -2,14 | 0,03 |
| Emitter Species * Number of Callers | -0,21 | 0,24 | -0,86 | 0,39 |

##### Hurdle 2

Occurrence

| | Estimate | SE | z | p |
|---|---|---|---|---|
| (Intercept) | -0,94 | 0,17 | -5,51 | <0,0001 |
| Emitter Species | 1,18 | 0,23 | 5.23 | <0,0001 |
| Number of Callers | 0,69 | 0,16 | 4,34 | <0,0001 |

Intensity

| | Estimate | SE | z | p |
|---|---|---|---|---|
| (Intercept) | 1 | 0,11 | 8,92 | <0,0001 |
| Emitter Species | -0,63 | 0,17 | -3,71 | 0,0002 |
| Number of Callers | -0,27 | 0,13 | -2,14 | 0,03 |
| Emitter Species * Number of Callers | -0,21 | 0,24 | -0,86 | 0,39 |

##### Hurdle 3

Occurrence

| | Estimate | SE | z | p |
|---|---|---|---|---|
| (Intercept) | -0,92 | 0,17 | -5,42 | <0,0001 |
| Emitter Species | 1,17 | 0,22 | 5.16 | <0,0001 |
| Number of Callers | 0,46 | 0,23 | 2,03 | 0,04 |
| Emitter Species * Number of Callers | 0,43 | 0,31 | 1,36 | 0,17 |

Intensity

| | Estimate | SE | z | P |
|---|---|---|---|---|
| (Intercept) | 0,99 | 0,11 | 8.81 | <0,0001 |
| Emitter Species | -0,58 | 0,16 | -3,67 | 0,0002 |
| Number of Callers | -0,33 | 0,11 | -3,06 | 0,002 |

| COMMUNITY | | | | |
|---|---|---|---|---|
| **B. Spring** | | | | |
| **Hurdle 5** | | | | |
| **Occurrence** | | | | |
| | **Estimate** | **SE** | **z** | **p** |
| (Intercept) | 0,05 | 0,16 | 0,30 | 0,77 |
| Emitter Species | 0,59 | 0,22 | 2,76 | 0,006 |
| Number of Callers | 0,97 | 0,22 | 4,43 | <0,0001 |
| Emitter Species * Number of Callers | -0,90 | 0,31 | -2,96 | 0,003 |
| **Intensity** | | | | |
| | **Estimate** | **SE** | **z** | **P** |
| (Intercept) | -0,08 | 0,38 | -0,20 | 0,84 |
| Emitter Species | -5.00 | 43.07 | -0,12 | 0,91 |
| Number of Callers | -0,54 | 0,35 | -1,53 | 0,13 |
| Emitter Species * Number of Callers | -4.17 | 60.81 | -0,07 | 0,95 |
| **Hurdle 6** | | | | |
| **Occurrence** | | | | |
| | **Estimate** | **SE** | **z** | **p** |
| (Intercept) | 0,04 | 0,17 | 0,25 | 0,80 |
| Emitter Species | 0,60 | 0,25 | 2,42 | 0,02 |
| Number of Callers | 0,52 | 0,18 | 2,93 | 0,003 |
| **Intensity** | | | | |
| | **Estimate** | **SE** | **z** | **P** |
| (Intercept) | -0,08 | 0,38 | -0,20 | 0,84 |
| Emitter Species | -10,07 | 6807.09 | -0,002 | 1,00 |
| Number of Callers | -0,54 | 0,35 | -1,53 | 0,13 |
| Emitter Species * Number of Callers | -11.32 | 9626.68 | -0,001 | 1,00 |
| **Hurdle 8** | | | | |
| **Occurrence** | | | | |
| | **Estimate** | **SE** | **z** | **p** |
| (Intercept) | 0,04 | 0,17 | 0,25 | 0,80 |
| Emitter Species | 0,60 | 0,25 | 2,42 | 0,02 |
| Number of Callers | 0,52 | 0,18 | 2,93 | 0,003 |
| **Intensity** | | | | |
| | **Estimate** | **SE** | **z** | **P** |
| (Intercept) | -0,08 | 0,44 | -0,19 | 0,85 |
| Emitter Species | -2.30 | 2.79 | -0,83 | 0,41 |
| Number of Callers | -0,55 | 0,36 | -1,54 | 0,12 |

| COAL | | | | |
|---|---|---|---|---|
| **C. Winter** | | | | |
| **Hurdle 9** | | | | |
| **Occurrence** | | | | |
| | **Estimate** | **SE** | **z** | **p** |
| (Intercept) | 0,52 | 0,16 | 3,30 | < 0.0001 |
| Emitter Species | 1,35 | 0,27 | 4,96 | < 0.0001 |
| Number of Callers | 0,50 | 0,21 | 2,33 | 0,02 |
| Emitter Species * Number of Callers | 0,38 | 0,38 | 1,02 | 0,31 |
| **Intensity** | | | | |
| | **Estimate** | **SE** | **z** | **p** |
| (Intercept) | 0,37 | 0,12 | 2,99 | 0,003 |
| Emitter Species | -0,24 | 0,26 | -0,92 | 0,36 |
| Number of Callers | -0,41 | 0,17 | -2,38 | 0,02 |
| Emitter Species * Number of Callers | 0,54 | 0,37 | 1,49 | 0,14 |
| **Hurdle 10** | | | | |
| **Occurrence** | | | | |
| | **Estimate** | **SE** | **z** | **p** |
| (Intercept) | 0,53 | 0,16 | 3,33 | 0,0009 |
| Emitter Species | 1,28 | 0,26 | 4,96 | < 0.0001 |
| Number of Callers | 0,63 | 0,18 | 3,57 | 0,0004 |
| **Intensity** | | | | |
| | **Estimate** | **SE** | **z** | **p** |
| (Intercept) | 0,37 | 0,12 | 2,99 | 0,003 |
| Emitter Species | -0,24 | 0,26 | -0,92 | 0,36 |
| Number of Callers | -0,41 | 0,17 | -2,38 | 0,02 |
| Emitter Species * Number of Callers | 0,54 | 0,37 | 1,49 | 0,14 |
| **Hurdle 11** | | | | |
| **Occurrence** | | | | |
| | **Estimate** | **SE** | **z** | **p** |
| (Intercept) | 0,52 | 0,16 | 3,30 | 0,001 |
| Emitter Species | 1,35 | 0,27 | 4,96 | < 0.0001 |
| Number of Callers | 0,50 | 0,21 | 2,33 | 0,02 |
| Emitter Species * Number of Callers | 0,38 | 0,38 | 1,02 | 0,31 |
| **Intensity** | | | | |
| | **Estimate** | **SE** | **z** | **p** |
| (Intercept) | 0,40 | 0,12 | 3,47 | 0,0005 |
| Emitter Species | -0,44 | 0,23 | -1,88 | 0,06 |
| Number of Callers | -0,31 | 0,15 | -2,03 | 0,04 |

### COAL

#### D. Spring

##### Hurdle 13

**Occurrence**

| | Estimate | SE | z | p |
|---|---|---|---|---|
| (Intercept) | 1,39 | 0,18 | 7,70 | < 0.0001 |
| Emitter Species | 0,88 | 0,30 | 2,89 | 0,004 |
| Number of Callers | 0,49 | 0,26 | 1,91 | 0,06 |
| Emitter Species * Number of Callers | -0,74 | 0,43 | -1,71 | 0,09 |

**Intensity**

| | Estimate | SE | z | p |
|---|---|---|---|---|
| (Intercept) | -0,42 | 0,36 | -1,15 | 0,25 |
| Emitter Species | -7,25 | 330,18 | -0,02 | 0,98 |
| Number of Callers | -0,36 | 0,36 | -0,99 | 0,32 |
| Emitter Species * Number of Callers | 9,99 | 466,95 | 0,02 | 0,98 |

##### Hurdle 14

**Occurrence**

| | Estimate | SE | z | p |
|---|---|---|---|---|
| (Intercept) | 1,36 | 0,18 | 7,74 | < 0.0001 |
| Emitter Species | 0,90 | 0,30 | 3,02 | 0,003 |
| Number of Callers | 0,23 | 0,20 | 1,13 | 0,26 |

**Intensity**

| | Estimate | SE | z | p |
|---|---|---|---|---|
| (Intercept) | -0,42 | 0,36 | -1,14 | 0,25 |
| Emitter Species | -5,96 | 91,20 | -0,07 | 0,95 |
| Number of Callers | -0,36 | 0,36 | -0,99 | 0,32 |
| Emitter Species * Number of Callers | 8,17 | 128,98 | 0,06 | 0,95 |

##### Hurdle 15

**Occurrence**

| | Estimate | SE | z | p |
|---|---|---|---|---|
| (Intercept) | 1,39 | 0,18 | 7,70 | < 0.0001 |
| Emitter Species | 0,88 | 0,30 | 2,89 | 0,004 |
| Number of Callers | 0,49 | 0,26 | 1,91 | 0,06 |
| Emitter Species * Number of Callers | -0,74 | 0,43 | -1,71 | 0,09 |

**Intensity**

| | Estimate | SE | z | p |
|---|---|---|---|---|
| (Intercept) | -0,40 | 0,36 | -1,11 | 0,27 |
| Emitter Species | -0,97 | 0,63 | -1,53 | 0,13 |
| Number of Callers | -0,13 | 0,31 | -0,41 | 0,68 |

### CRESTED

#### E. Winter

##### Hurdle 17

**Occurrence**

| | Estimate | SE | z | p |
|---|---|---|---|---|
| (Intercept) | 0,51 | 0,15 | 3,51 | 0,0005 |
| Emitter Species | 1,18 | 0,26 | 4,53 | <0,0001 |
| Number of Callers | 0,21 | 0,21 | 1,02 | 0,31 |
| Emitter Species * Number of Callers | 0,85 | 0,37 | 2,30 | 0,02 |

**Intensity**

| | Estimate | SE | z | p |
|---|---|---|---|---|
| (Intercept) | 0,36 | 0,12 | 3,15 | 0,002 |
| Emitter Species | -0,68 | 0,38 | -1,81 | 0,07 |
| Number of Callers | -0,23 | 0,16 | -1,39 | 0,16 |
| Emitter Species * Number of Callers | -0,41 | 0,53 | -0,77 | 0,44 |

##### Hurdle 18

**Occurrence**

| | Estimate | SE | z | p |
|---|---|---|---|---|
| (Intercept) | 0,53 | 0,15 | 3,55 | 0,0004 |
| Emitter Species | 1,03 | 0,24 | 4,33 | <0,0001 |
| Number of Callers | 0,51 | 0,17 | 3,06 | 0,0020 |

**Intensity**

| | Estimate | SE | z | p |
|---|---|---|---|---|
| (Intercept) | 0,36 | 0,12 | 3,15 | 0,002 |
| Emitter Species | -0,68 | 0,38 | -1,81 | 0,07 |
| Number of Callers | -0,23 | 0,16 | -1,39 | 0,16 |
| Emitter Species * Number of Callers | -0,41 | 0,53 | -0,77 | 0,44 |

##### Hurdle 19

**Occurrence**

| | Estimate | SE | z | p |
|---|---|---|---|---|
| (Intercept) | 0,51 | 0,15 | 3,51 | 0,0005 |
| Emitter Species | 1,18 | 0,26 | 4,53 | <0,0001 |
| Number of Callers | 0,21 | 0,21 | 1,02 | 0,31 |
| Emitter Species * Number of Callers | 0,85 | 0,37 | 2,30 | 0,02 |

**Intensity**

| | Estimate | SE | z | p |
|---|---|---|---|---|
| (Intercept) | 0,35 | 0,12 | 3,04 | 0,002 |
| Emitter Species | -0,47 | 0,24 | -2,01 | 0,04 |
| Number of Callers | -0,28 | 0,15 | -1,80 | 0,07 |

| CRESTED | | | | |
|---|---|---|---|---|
| **F. Spring** | | | | |
| **Hurdle 21** | | | | |
| **Occurrence** | | | | |
| | **Estimate** | **SE** | **z** | **p** |
| (Intercept) | 3,09 | 0,52 | 5,95 | <0,0001 |
| Emitter Species | -0,56 | 0,59 | -0,95 | 0,34 |
| Number of Callers | 2,13 | 0,74 | 2,90 | 0,004 |
| Emitter Species * Number of Callers | -1,82 | 0,83 | -2,19 | 0,03 |
| **Intensity** | | | | |
| | **Estimate** | **SE** | **z** | **p** |
| (Intercept) | 0,01 | 0,50 | 0,02 | 0,99 |
| Emitter Species | -1,35 | 0,85 | -1,59 | 0,11 |
| Number of Callers | 0,65 | 0,71 | 0,92 | 0,36 |
| Emitter Species * Number of Callers | -0,37 | 1,21 | -0,31 | 0,76 |
| **Hurdle 22** | | | | |
| **Occurrence** | | | | |
| | **Estimate** | **SE** | **z** | **p** |
| (Intercept) | 2,51 | 0,28 | 9,06 | < 0,0001 |
| Emitter Species | 0,20 | 0,37 | 0,55 | 0,58 |
| Number of Callers | 1,00 | 0,31 | 3,23 | 0,001 |
| **Intensity** | | | | |
| | **Estimate** | **SE** | **z** | **p** |
| (Intercept) | 0,01 | 0,50 | 0,02 | 0,99 |
| Emitter Species | -1,35 | 0,85 | -1,59 | 0,11 |
| Number of Callers | 0,65 | 0,71 | 0,92 | 0,36 |
| Emitter Species * Number of Callers | -0,37 | 1,21 | -0,31 | 0,76 |
| **Hurdle 24** | | | | |
| **Occurrence** | | | | |
| | **Estimate** | **SE** | **z** | **p** |
| (Intercept) | 2,51 | 0,28 | 9,06 | < 0,0001 |
| Emitter Species | 0,20 | 0,37 | 0,55 | 0,58 |
| Number of Callers | 1,00 | 0,31 | 3,23 | 0,001 |
| Emitter Species * Number of Callers | | | | |
| **Intensity** | | | | |
| | **Estimate** | **SE** | **z** | **p** |
| (Intercept) | -0,06 | 0,48 | -0,12 | 0,90 |
| Emitter Species | -1,30 | 0,88 | -1,48 | 0,14 |
| Number of Callers | 0,51 | 0,61 | 0,84 | 0,40 |
